# Cell type catalog of middle turbinate epithelium

**DOI:** 10.1101/2022.10.09.511503

**Authors:** Fasil Mathews, Victoria Sook Keng Tung, Robert Foronjy, Marina Boruk, James A Knowles, Oleg V Evgrafov

## Abstract

**Importance:** Elucidation of the cellular makeup of the middle turbinate provides a foundation for future studies of pathogenesis of sinonasal disease. Neural progenitors and pluripotent basal cells found in middle turbinate mucosa potentially can be used to develop cellular models to study brain disorders or in regenerative medicine to substitute neuronal tissues.

**Objective:** Single cell RNA-sequencing (scRNA-seq) of middle turbinate mucosa was performed to create the first single cell transcriptome catalog of this part of the human body.

**Design:** Samples were obtained from the head of the middle turbinate from healthy volunteers. After the specimen was prepared per lab protocol, cells were dissociated, suspended, and counted. Single cell libraries were then prepared according to the 10x Genomics protocol and sequenced using NovaSeq 6000 (Illumina). Sequencing data were processed using Cell Ranger, and clustering and gene expression analysis was performed using Seurat. Cell types were annotated using known markers and data from other single cell studies.

**Setting:** Single center, tertiary care center

**Participants:** Healthy volunteer

**Intervention(s) (for clinical trials) or Exposure(s) (for observational studies):** None

**Main Outcome(s) and Measure(s):** Identification of cell types of middle turbinate mucosa through expression profiling of single cells using known markers

**Results:** 14 unique cell types were identified, including serous, goblet, club, basal, ciliated, endothelial, and neural progenitor cells, as well as multiple types of blood cells.

**Conclusions and Relevance:** This catalog provides a comprehensive depiction of the cellular composition of middle turbinate mucosa. By uncovering the cellular stratification of gene expression profiles in healthy middle turbinate epithelium, the groundwork has been laid for further investigation into the molecular pathogenesis and targeted therapy of sinonasal disease.

**Key Points:** *Question:* What is the cellular makeup of human middle turbinate mucosa?

*Findings:* Single-cell RNA sequencing revealed 14 cell types in middle turbinate epithelium, including neural progenitors, a previously unrecognized component of middle turbinate epithelium.

*Meaning:* Gene expression profiles of middle turbinate mucosa cell types are concordant with other respiratory mucosa transcriptomic data with notable heterogeneity in serous and basal cells.□

## Introduction

The function of an organ is defined by the coordinated actions of different cell types. Until recently, cell types could only be differentiated by cell morphology, functional properties and, most typically, by marker proteins. Nevertheless, some cell types could not be distinguished from each other because the expression levels of proteins defining the difference was not known. Single cell RNA-sequencing (scRNA-seq) allows for whole transcriptome analysis, thus removing bias from analysis of expression profiles of different cell types. Cells are grouped based on similarity of their expression profiles using cluster analysis, and “marker” genes with statistically significant differences between cells in one cluster and cells in all other clusters are then used to identify cell types.^1,2^ This approach has already demonstrated its superiority compared to classical identification of cell types.^3–5^

In this study, we performed scRNA-seq of a mucosal biopsy from the anterior portion of the middle turbinate. Nasal turbinates are bowed shelves projecting from the lateral nasal wall and base of skull; they function to warm and humidify inspired air by regulating nasal airflow. Additionally, as the first structures of the upper airway to receive inspired air, they play a role in immune surveillance.^6–8^ They have been described to be composed of columnar pseudostratified respiratory epithelium containing basal cells, ciliated cells, glandular epithelial cells, and olfactory epithelial cells.^9–11^ Many of these studies looked at nasal cavity tissue or turbinate tissue in general; however, the specific cellular framework of the middle turbinate has not yet been described.

As more targeted molecular approaches to the management of chronic sinonasal disease are being explored, we believe better elucidation of the cellular makeup of the middle turbinate will provide a foundation for future research in the pathogenesis and medical treatment of sinonasal disease. Furthermore, multiple studies have also shown that middle turbinate epithelium contains pluripotent cells which could used both in research and regenerative medicine.^12–15^ It can also be used to produce neural progenitor cell cultures, a useful cellular model of several disorders of the brain.^16^

## Methodology

### Biopsy collection and sample preparation

Biopsies were obtained from patients without any history of sinonasal disease or surgery or immunocompromise. Tissue samples were obtained from the superior-medial region of the head of the middle turbinate; laterality was determined by ease of access. The mucosa was anesthetized and decongested with topical 1% lidocaine and 0.05% oxymetazoline. After 5 minutes, 0.3 ml of 1% lidocaine and 1:100,000 epinephrine was injected into the targeted mucosal site under visualization. 2mm cupped forceps were used to obtain biopsy specimens. These samples were immediately transported to the lab in Leibovitz’s L-15 Medium (ThermoFisher) supplemented with Antibiotic-Antimycotic solution (ThermoFisher) and processed to prepare single cell suspension. Biopsy pieces were minced with two scalpels in Petri dish with small amount of cold Hank’s buffer without magnesium or calcium to prevent drying and then transferred to a 1.5 ml Eppendorf tube and washed twice with 1 ml cold Hank’s buffer. Cells were dissociated in 250 µl of 0.25% Trypsin-EDTA at 37ºC while shaking at 500 rpm. After 10 minutes, the suspension was mixed by pipetting and returned to thermomixer for another 10 minutes. After another mixing, the cells were sedimented by centrifugation at 300 relative centrifugal force for 5 minutes, resuspended in 500 µl of cold Hank’s solution with BSA, filtered using 40 µm FLOWMI cell strainer cell suspension into a 1.5 ml Eppendorf tube, centrifuged again, and finally resuspended in 50 µl of Hank’s with BSA. The concentration and viability of cells was determined using a hemocytometer and Trypan blue. After counting, single cell libraries were prepared according to the 10x Genomics protocol CG000183 on Chromium controller (10x Genomics).

### Data analysis

Sequencing data was processed using bcl2fastq2 v2.20 to convert bcl files generated by NovaSeq 6000 sequencing instrument to fastq files and perform simultaneous demultiplexing. Fastq files were processed using TrimGalore v. 0.6.5 to automate quality and adapter trimming and perform quality control. Fastq files were analyzed using Cell Ranger v. 6.1.2 (10x Genomics) and the ‘count’ command was used to generate raw gene-barcode matrices aligned to the GRCh38 Ensembl v93-annotated genome. The data were further analyzed using Seurat v.4.10 to filter cells with low-quality data, normalize and scale gene expression, and cluster cells by a graph-based algorithm.^17^ We performed clustering using different values of *resolution* parameters to assess relationships between groups of cells. Furthermore, we annotated clusters using known markers and data from other single-cell studies of relevant tissues or cell types. For visualization purposes, we generated UMAP graphs using algorithms implemented in Seurat.

## Results

Analysis of 541,837,594 transcripts from 21,565 cells (mean reads per cell: 25,126) identified substantial heterogeneity of gene expression profiles and clear tendency to cluster according to cell types. Those genes expressed at a higher level than in other clusters, and especially those expressed at a high level in the cluster, were considered as marker genes and used for annotation of cells in the cluster.

14 cell types were identified using previously described cell-type specific markers (Figure 1A). These included cell types that typically compose respiratory epithelium, such as serous, goblet, club, basal, ciliated, endothelial, and neural progenitor cells (Figure 1B), as well as multiple types of blood cells (Figure 1C). Additionally, we identified a group of likely apoptotic cells as well as a group of cells without any characteristic known markers of any cell type.

**Figure 1.**
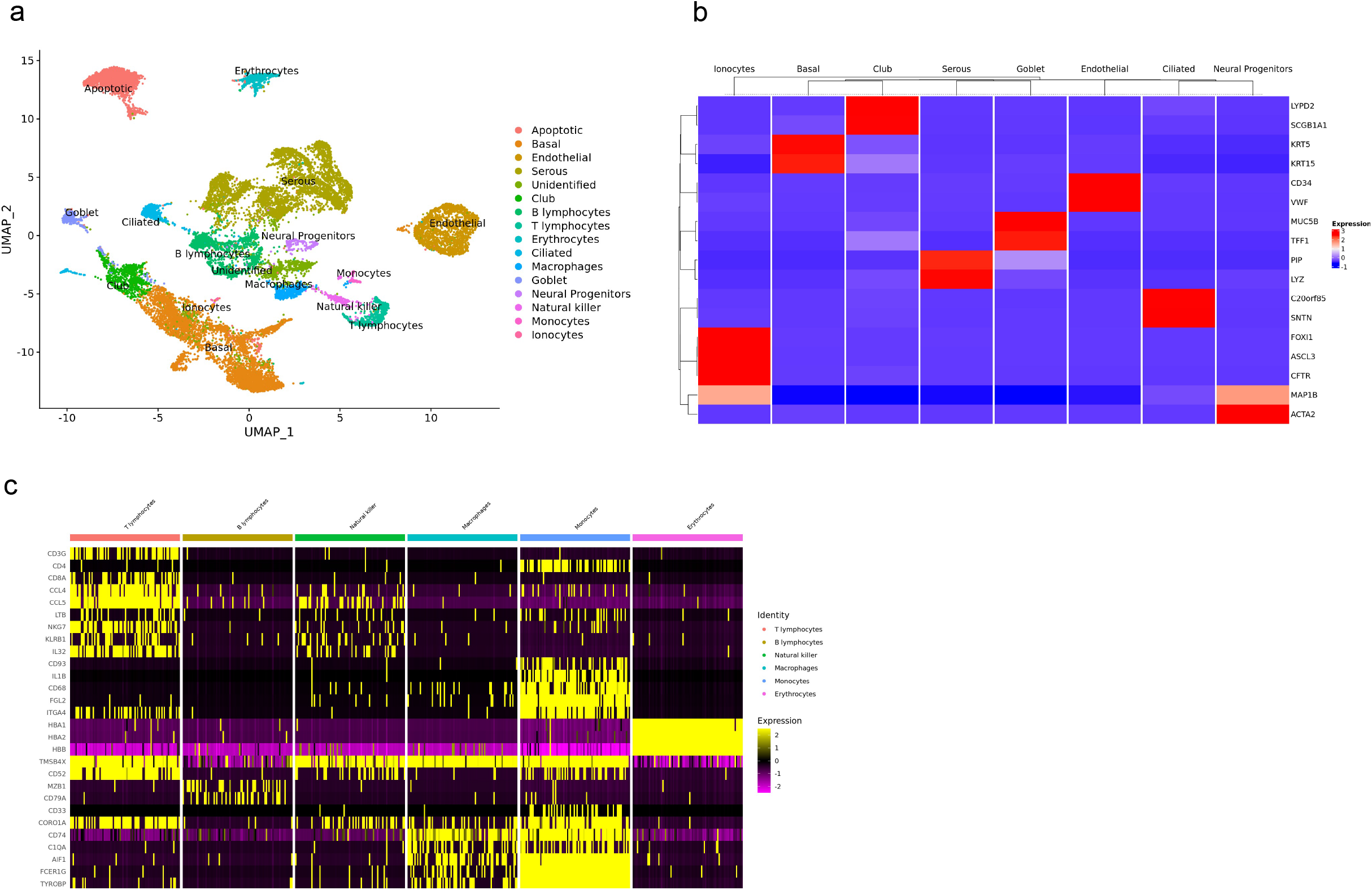
Aggregate analysis of cells from human middle turbinate epithelium. (A) UMAP dimensionality reduction plot of 21,565 human middle turbinate cells from sample SEP310 (healthy donor). Clusters associated with cell types according to known markers. (B) Heatmap using average expression of main respiratory epithelium cell types with marker genes for specific cell types. (C) Heatmap using average expression of blood cell types with marker genes for specific cell types.

### Glandular Epithelial Cells

#### Serous cells

Serous cells secrete zymogens, enzymes, and antibodies to defend airways against bacteria and viruses; these include lysozyme, lactoferrin, zymogen granule protein, and bactericidal permeability-increasing protein. We identified a large cluster of serous cells characterized by high expression of *LYZ, STATH, LTF, ZG16B, BPIFA1, BPIFB1, S100A1* and *PIP* (Figure 2)^18–31^; the latter two play a role in regulation of calcium content in the secretions.^18,20,32^

**Figure 2.**
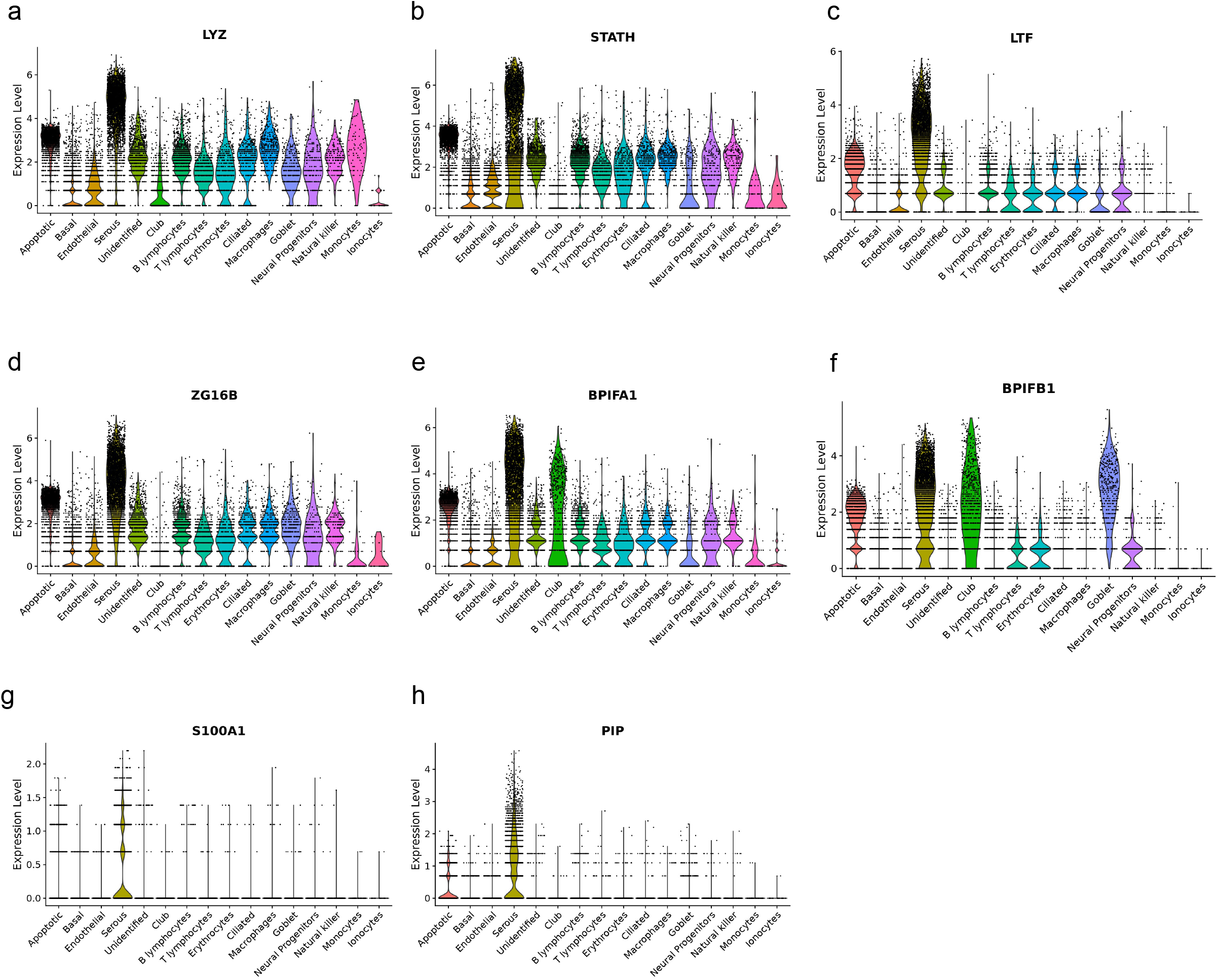
Expression of marker genes of serous cells in different cell types. Violin plots show the expression distribution of various markers of serous cells (A-H). Each dot represents a single cell

#### Goblet Cells

*SPDEF, MUC5B, MUC5AC, FOXA3, TFF1* and *TFF3* are known to be expressed in mucus-producing goblet cells of human respiratory epithelia.^33–36^ *BPIFB2* has also been shown to be highly expressed in goblet cells in pig submucosal glands.^37,38^ We found a cluster characterized by expression of these genes (Figure 3), which identified this cluster as goblet cells. Some of these genes (*MUC5B, TFF1*, and *TFF3)* were previously shown to be over-expressed in middle turbinate samples from patients with CRS.^39,40^

**Figure 3.**
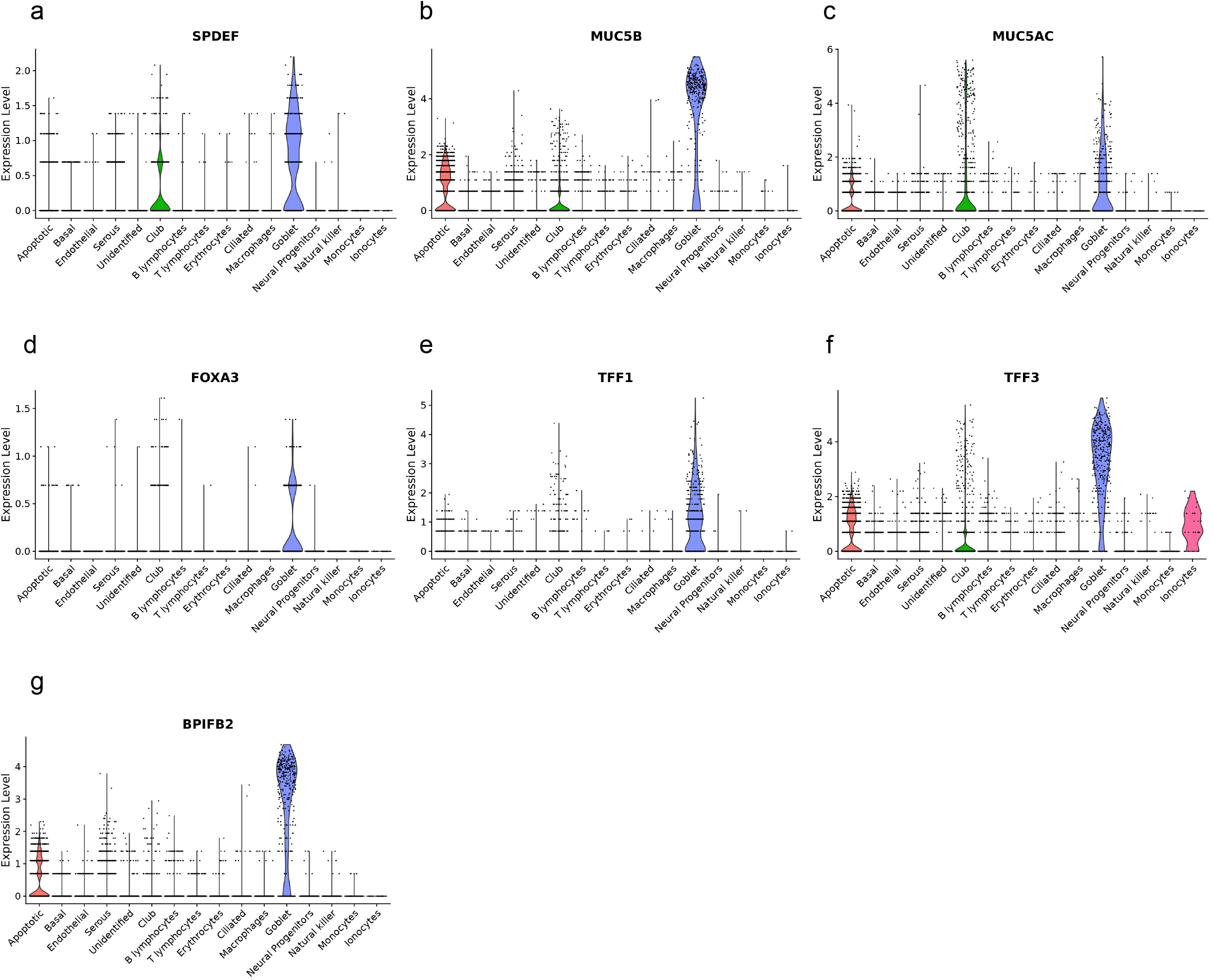
Expression of marker genes of goblet cells in different cell types. Violin plots show the expression distribution of various markers of goblet cells (A-G). Each dot represents a single cell.

#### Club Cells

Club cells were identified by expression of known marker *SCGB1A1*. We found that club cells were also characterized by exclusive expressions of *LYPD2* and *S100P* (Figure 4). *LYPD2* and *S100P* have both been detected in human nasal respiratory epithelium in other studies but have not been previously localized to club cells.^41–43^ Given these genes were highly expressed by cells in this cluster, we suggest they may be used as markers for turbinate club cells. Although Hogan and Tata suggested *SCGB3A2* is a marker of club cells in mice, we do not have data to support it is in human middle turbinate.^44^ Of note, Deprez et al. was able to identify deuterosomal cells, or precursors to ciliated cells, in airway epithelium^43^; however, differential expression of these marker genes (*DEUP1, FOXN4, CDC20B)* was not detected in our study.

**Figure 4.**
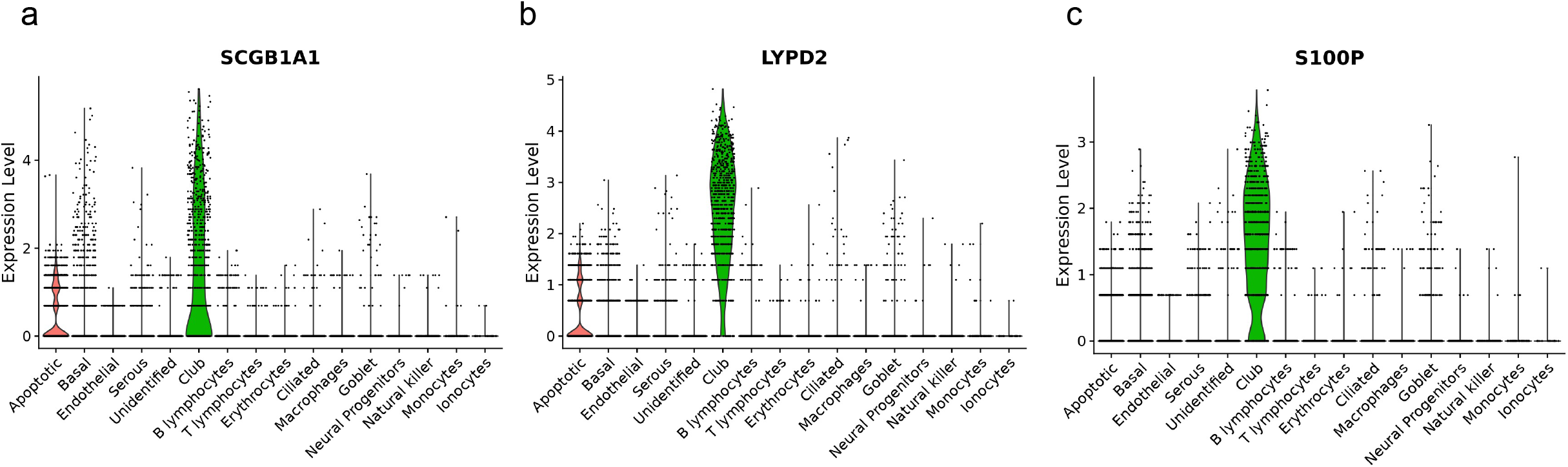
Expression of marker genes of club cells in different cell types. Violin plots show the expression distribution of various markers of club cells (A-C). Each dot represents a single cell.

### Non-Secretory Epithelial Cells

#### Basal Cells

We identified a large cluster of respiratory epithelial basal cells in the middle turbinate. *KRT5* is usually used to define basal cells in respiratory epithelium; in our sample, *KRT15* was also a useful marker (Figure 1B).^45–47^.

#### Ciliated Cells

Ciliated nasal respiratory epithelial cells had a distinctive molecular profile (Figure 1B). Sentan (*SNTN)* is an identifiable component of the apical structure of vertebrate motile cilia.^48^ Chromosome 20 open reading frame 85, which codes for Low in Lung Cancer 1 (LLC1) protein, has also been localized in the cilia of normal lung epithelium, as well as in an unspecified location of nasal mucosa.^49^

#### Ionocytes

Pulmonary ionocytes are cells rich in the CFTR chloride channel. These cells were detected in our sample as having high expression of *CFTR, ASCL3*, and *FOXI1* differentiating them from other cell types (Figure 1B).^33,43^

### Blood Cells

In addition to epithelial cells, several types of white blood cells were identified. B-cells were identified by expression of *MZB1* (Figure 1C). T-lymphocytes expressed *CD3, CD4*, and *CD8* along with *CCL4, CCL5, LTB*, and *NKG7*. ^50–52^ Natural killer cells largely did not expreses *CD3* and a few cells expressed *CD4*. They also expressed *KLRB1, IL32*, and *CORO1A* (Figure 1C).^53–55^

Macrophages expressed *CD74* but not *CD3*. They also expressed *C1QA, AIF1, FCER1G*, and *TYROBP*.^56^ Monocytes expressed several markers of macrophages or monocytes, including *IL1B, CD68*, and *FGL2*; however, they were specified as monocytes as they expressed *CD33* and *CD93*.^57,58^ It is interesting that this group also expressed multiple markers of neutrophils, including *FCGR3A, ICAM1, ITGAM, IL17RA*, and *ITGA4*. As such, this cluster may also contain some neutrophils. Lastly, erythrocytes were identified by expression of *HBA1, HBA2*, and *HBB* (Figure 1C).^59^

### Other Cell Types

Other annotated cell types included neural progenitor cells and endothelial cells. Previously we used superior turbinate and middle turbinate biopsies to produce cell lines of neural progenitor cells known as cultured neural progenitors derived from olfactory neuroepithelium (CNON)^16^, and we identified in this study a cell type with an expression profile very close to CNON (expressing *ACTA2, MAP1B*, and *COL1A2)* thus being likely a precursor of CNON (Supplemental Figure 1). Vascular endothelial cells were characterized by expression of *ACKR1, SELE, SOX18, MME, EDN1, PLVAP* and *VWF* (Supplemental Figure 2).^60^

Cell types from two clusters were not annotated as a specific cell type. One of the clusters was distinguished from others by substantially higher expression of mitochondrial genes, a characteristic of apoptotic cells (Supplemental Figure 3).^61–63^ Notably, gene characteristic of club cells (*LYPD2* and *SCGB1A1)* and serous cells (*DMBT1*) were also expressed in this cluster albeit at a substantially lower level, suggesting these cell types contributed to this apoptotic cluster. For another cluster, we were not able to identify any marker genes. Overall, this cluster was characterized by high expression of ribosomal protein genes (78 out of 130 differentially expressed genes), indicating increased protein production. This group may represent a different functional state of a previously identified cell type or a unique cell type.

## Discussion

We have generated the first single-cell catalog of normal human middle turbinate epithelium, revealing 14 unique cell types. During this workflow, we encountered significant heterogeneity of serous and basal cells that warrants further discussion.

We broadly categorized 5 serous clusters into 2 groups (Figure 5A-B). Although several of the marker genes aforementioned showed a similar expression pattern in all subgroups (Supplemental Figure 4), only Serous1-3 expressed *DMBT1* at a high level (Figure 5C); this protein is a known marker of serous cells^37^ and has been identified in serous acini cells in human submandibular and labial tissue.^21^ In comparison, Serous4 and Serous5 had higher expression of various immunoglobulins (Figure 5D-F). Although both are involved in the immune response, it is clear there is variation in the protein composition of secretions from serous cells in the middle turbinate. We were surprised to find that these two groups were very different in the overall level of gene expression and correspondingly in the number of expressed (detectable) genes (Figure 6). This is unlikely an artifact of the analysis parameters as some serous marker genes as well as housekeeping genes were expressed at a similar level in both groups, including housekeeping genes. This is unlikely an artifact of the analysis parameters as some serous marker genes, as well as housekeeping genes, were expressed at a similar level in both groups.

**Figure 5.**
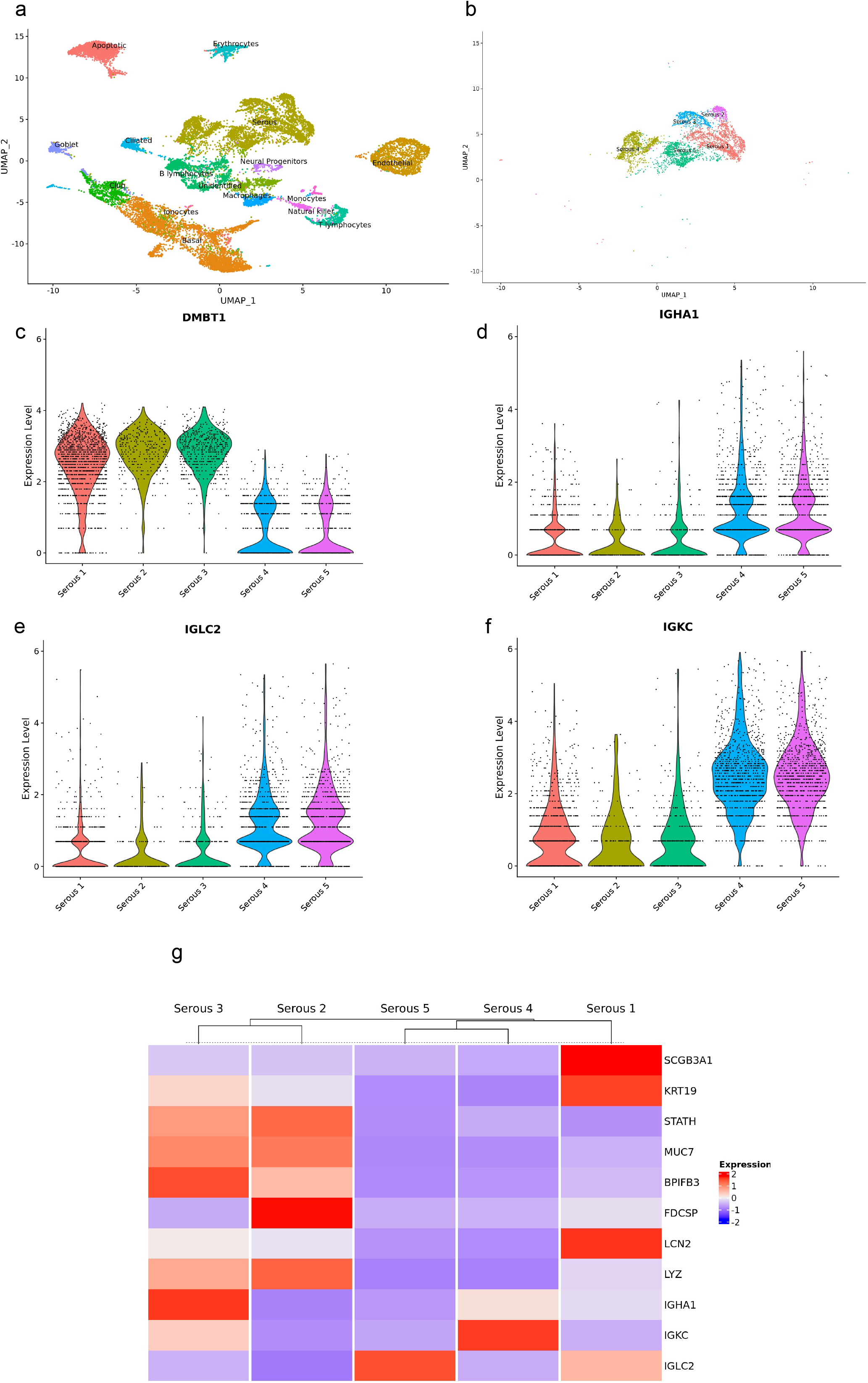
Expression of genes differentiating serous cell subgroups. (A) UMAP dimensionality reduction plot of 21,565 human middle turbinate cells from sample SEP310 (healthy donor). Clusters associated with cell types according to known markers. (B) UMAP dimensionality reduction plot of serous cell subgroups. Violin plots show elevated expression of *DMBT1* in Serous1-3 (C) and various immunoglobulins in Serous4 and Serous5 (D-F) with each dot representing a single cell. (G) Heatmap showing average gene expression in cell subgroups depicts selected gene expression among serous cell subgroups. Serous1 cluster had higher expression of *SCGB3A1, KRT19* and lower expression of *STATH, MUC7, BPIFB3* than Serous2 and Serous3.

We did find genes differentiating all five subgroups (Figure 5G). *FDCSP* was highly expressed in Serous2 cluster. It is known to play a role in immune response, but the significance of this variation between groups is unclear. *STATH* has been identified in sinonasal secretions and was noted to be expressed at higher levels in patients with CRS;^31^ expression levels of this gene distinguished Serous2-4 from Serous1 and Serous5. It is unclear if these subgroups represent different inflammatory states of the same cell type or truly different cell subtypes. Lastly, a study of submucosal glands in pigs was able to differentiate serous acinar cells from serous ductal cells; in accordance with their data, *LCN2* was not detected in Serous4 cluster suggesting this subgroup consisted of only acinar (Figure 5E). Taken together, these data suggest serous cells are not a uniform group of cells, and this heterogeneity highlights the complex role serous cells play in innate immune function and maintaining homeostasis of sinonasal secretions.

Basal cells are often characterized by expression of *TP63* or *DLK2*.^35,44,64,65^ Interestingly, Deprez et al. had grouped all basal cells in olfactory epithelium as *DLK2*-high;^43^ however, in our data, these two genes are expressed (and show nearly identical expression patterns) only in Basal2 and Basal3 clusters (Supplemental Figure 5). Goldstein et al. found that *SERPINB3* colocalized to *KRT5+* basal cells in respiratory epithelium that lacked olfactory neurons; in congruence with their data, our analysis revealed *SERPINB3+* subgroups (Basal1 and Basal4) also expressed other known respiratory epithelial genes including *HES4* and *MUC1* (Supplemental Figure 5).^36^ Additionally, *SERPINB3-* basal cell subgroups (Basal2 and Basal3) had elevated expression of *SPINK5*, which localized to olfactory basal cells.^36^ These data suggest *SERPINB3* (along with the aforementioned correlated genes) could be used as a marker to differentiate respiratory basal cells from olfactory basal cells in the middle turbinate.

Basal4 cluster differentiated from Basal1 (and all other clusters) by expression of genes involved in cell proliferation and division including *MKI67* and *TOP2A* (Supplemental Figure 5).^43^ Based on similarity of expression profiles, we suggest these cells are from Basal1 cluster entering into a proliferative state; nevertheless, there are genes expressed in Basal4, but not in Basal1 cluster, which are not related to cell proliferation (*KRT15, KRT17)*.

We would like to highlight the first description of club cells in the nasal turbinates. Club cells and their role in chronic obstructive pulmonary disease (COPD) has been extensively described.^66–68^ A recent study showed that almost ¼ of patients with COPD had comorbid CRS.^69^ Given the similarities in pathogenesis, namely chronic inflammation and hypersecretion of mucous, club cells may be a potential link in future areas of research on disease development and treatment. Of note, Deprez et al. had previously grouped club and goblet cells together in human respiratory epithelium^43^; however, our data shows sufficient differentiation between these cell types in human middle turbinate.

Ionocytes were only recently described in airway epithelium, and this is first study describing them in the nasal turbinate. Our data may assist in further drug development in cystic fibrosis and other mucociliary disorders by improving localization of the CFTR channel.

Although olfactory epithelium has been inconsistently reported to be found in the middle turbinate, we did not detect any olfactory cells, likely as our biopsy was from the anterior rather than the superomedial portion.^7,70^ Additionally, mRNA of tuft/brush cell marker genes was either not found or only a few molecules were detected in the whole cell population.^71–73^

Lastly, we compared this single cell data which was collected from a healthy individual (SEP310) with data from a sample from patient with schizophrenia (SEP313). Both samples were processed at the same time. Using reference mapping, gene expression data of the compared sample was presented in the same coordinates as the reference sample, and cell types were identified in accordance with our prior annotation of “reference” datasets (“label transfer”). A similar analysis was performed on the scRNA-seq data of olfactory epithelium from “Patient 4” in Durante et al.^74^ Both analyses demonstrated broadly similar results, with blood cells being expectedly the most variable component (Supplemental Figure 6). Apoptotic cells in SEP313 were much lower in number, but their expression profile matched that of apoptotic cells in SEP310; while, most of the apoptotic cells from olfactory epithelium showed a different expression profile, suggesting they originated from different cell types. Both samples demonstrated heterogeneity of basal and serous cells, as well as identified neural progenitor cells in substantial numbers.

There are several limitations to our methodology to concede. Our analysis was performed from a small portion of mucosa derived from the head of the middle turbinate. Some cell types may have been missed due to insufficient sample size. Furthermore, the cell type composition from the head of the middle turbinate may not be generalizable to the middle or tail of the turbinate. Lastly, any local changes in the tissue due to patient factors or as a result of the biopsy may have resulted in minor changes in the cell type composition. The significance of our findings can be strengthened with corroboration with histologic analysis of turbinate epithelium and proteomic analysis of mucin and other secretions.

## Conclusion

This single cell reference map provides the first depiction of the cellular composition of middle turbinate mucosa. We identified cell types previously described in respiratory epithelium, such as basal, club, goblet and serous cells as well as potential neural progenitors. We described substantial heterogeneity in expression profiles and overall level of gene expression in basal and serous cells, suggesting the existence of distinct subtypes with different biological roles. Future areas of investigation include spatial transcriptomics analysis to localize cell types to specific anatomic regions of the middle turbinate.

---

Supplemental Figure 1

Title: Expression of marker genes of neural progenitor cells in different cell types.

Legend: Violin plots show the expression distribution of various markers of cultured neural progenitors derived from olfactory neuroepithelium (CNON) (A-C). Each dot represents a single cell.

Supplemental Figure 2

Title: Expression of marker genes of endothelial cells in different cell types.

Legend: Violin plots show the expression distribution of various markers of endothelial cells (A-G). Each dot represents a single cell.

Supplemental Figure 3

Title: Heatmap showing expression of mitochondrial genes in different cell types.

Legend: Heatmap using average expression of cell types with mitochondrial genes.

Supplemental Figure 4

Title: Expression levels of serous cell subgroups

Legend: Violin plots show elevated levels of number of expressed (detectable) genes, total number of reads and overall gene expression (sum of normalized expression of all genes) in Serous1-3 compared to Serous 4 and Serous 5 subgroups. Serous marker genes (*ZG16B, LYZ*) and housekeeping genes (*ACTB, GADPH*) were expressed at a similar level in all groups.

Supplemental Figure 5

Title: Gene expression of basal cell subgroups

Legend: (A) UMAP dimensionality reduction plot of 21,565 human middle turbinate cells from sample SEP310 (healthy donor). Clusters associated with cell types according to known markers. (B) UMAP dimensionality reduction plot of basal cell subgroups. (C) Heatmap depicting selected gene expression among basal cell subgroups.

Supplemental Figure 6

Title: Comparative UMAP dimensionality reduction plots

Legend: Single cell data from middle turbinate (SEP310, (A)) used as a reference for other plots. (B) Single cell data from middle turbinate from patient with schizophrenia projected onto reference UMAP structure. (C) Single cell data from olfactory neuroepithelium from patient 4 downloaded from GSE139522 study and processed through our workflow projected onto reference UMAP structure.^74^

